# Zero-shot biological reasoning with open-weights large language models reproduces CRISPR screen based prediction of synthetic lethal interactions

**DOI:** 10.64898/2026.01.28.702211

**Authors:** Aurel Prosz, Zsofia Sztupinszki, Miklos Diossy, Oz Kilim, Bogumil Zimon, Istvan Csabai, Zoltan Szallasi

## Abstract

Identifying clinically relevant synthetic lethal interactions has great potential for uncovering novel therapeutic vulnerabilities in cancer. Current approaches rely on machine learning models that estimate probabilities of synthetic lethal interactions, without supplying explicit knowledge of the underlying biology and lack the human-readable interpretation leading to the prediction. Large Language Models (LLMs) represent a new class of tools capable of reasoning and leveraging extensive biological knowledge acquired from relevant literature during their pretraining. Here, we tested multiple open-weight LLMs for their ability to predict known and novel synthetic lethal interactions. We found that most of the tested models were better at reconstructing the results of three known genome-wide CRISPR knockout screens than random chance and non-LLM-based methods, and observed that their performance was related to the parameter-size of the model, and on average benefited little from additional pathway and genetic information apart from what they already possess when estimating the likelihood of a synthetic lethal relationship. After selecting the model that offered the best balance of predictive performance and computational efficiency for our use case (Qwen2.5-32B-Instruct, 0.715 AUROC), we performed an in silico screen of 398,277 gene pairs from 893 clinically relevant genes. Our goal was to highlight the potential of open-weights LLMs as scalable, context-aware prioritization tools for synthetic lethal interactions, and to lay the groundwork for predicting higher-order genetic interactions.

## Introduction

Synthetic lethality (SL) is a genetic interaction where the combined, simultaneous inactivation of two genes, by e.g. loss of function mutation or small molecule inhibitors, results in cell death, while the inactivation of either gene alone allows cell viability [1]. SL provides a principled strategy for uncovering cancer-specific vulnerabilities, exemplified by the clinical success of PARP inhibitors in tumors with defects in homologous recombination repair [2]. Advances in CRISPR-based genetic screens have expanded the catalogue of SL interactions across cell lines and genetic backgrounds [3]. These methods are powerful to identify novel synthetic interactions between the loss of a single gene and the pharmacological modification of a specific target such as *PARP* [4]. It is quite likely, however, that there might be more complex, multigenic SL interactions that could be therapeutically exploited [5]. For example, in lung cancer models co-occurrence of *KRAS* and *KEAP1* mutations causes a synthetic lethality with glutaminase inhibition [6]. While combinatorial CRISPR screening is possible, genome scale combinatorial screening for multigenic SL interactions is experimentally and financially not feasible when a combination of three or more genes is investigated [5, 7–9]. In silico approaches, if they accurately reflect the underlying biology, may provide a method to perform such combinatorial screening on a larger scale than “wet lab”-based screening currently allows [10–12].

Another major challenge in the translatability of novel SL predictions is their context specificity, which can be summarized as the problem with SL predictions depending on the exact lineage, genotype or even culture conditions [13, 14]. Partly, this has motivated broad, large scale functional mapping efforts, such as the Cancer Dependency Map, for mapping out the cell-line specific dependencies across hundreds of different cell lines derived from different cancer types [15].

Current state-of-the-art computational approaches to SL prediction are predominantly data-driven, that rely on supervised machine learning models that integrate protein–protein interaction networks, gene expression values, and pathway annotations [16, 17]. Graph neural networks, hybrid graph–transformer architectures and models based on knowledge-graph reasoning achieve high predictive capability on benchmark datasets [18, 19] with graph autoencoders emerging as state-of-the-art in recent comparisons as well [16]. However, these approaches tend to underperform when applied to under-annotated genes or unseen contexts and their mechanistic interpretability can be often highly limited, which suggests that learned predictions might reflect dataset-specific correlations rather than true biological reasoning [20]. Large Language Models (LLMs) represent a fundamentally different paradigm. Pretrained on vast corpora of biomedical, genomic and other related text, LLMs capture statistical patterns in language that reflect published associations among genes, pathways, and phenotypes. These representations may correlate with functional biology, but they should not be interpreted as direct or verified encoding of biological mechanisms. [21, 22]. Recent work has shown that LLM-derived gene embeddings can result in better gene–gene interaction and perturbation prediction, often even outperforming specialized models, particularly in cold-start settings [23, 24]. Independent benchmarks and studies have further demonstrated that LLMs can reason accurately about gene function and genetic interactions when appropriately prompted or used special scaffolding [24–28]. Despite the advances listed above, LLMs have not been directly applied to synthetic lethality prediction in a high-throughput way. Existing efforts are limited to explaining known SL interactions using specialized, fine-tuned models and external knowledge graphs, rather than predicting novel gene pairs *en masse*, with interpretations [18, 29]. Notably, recent comprehensive benchmarks of SL prediction methods include no pure LLM-based approaches, underscoring a methodological gap in the field [16].

## Results

As a practical benchmark and explorative study, we started by downloading already published genome-wide CRISPR knockout screens. As a first step, we systematically evaluated six open-weights language models (Llama3-OpenBioLLM-8B [30], Qwen2.5-7B-Instruct, Qwen2.5-32B-Instruct, Qwen2.5-72B-Instruct [31], gpt-oss-20b, gpt-oss-120b [32]) by testing their ability to reproduce results from a well-characterized Olaparib CRISPR screening experiment performed in *TP53* -deficient RPE1 cells [3]. We chose open-weights models, since making predictions on a scale of million gene pairs or higher order combinations require cost-effective, but also high-performing models. The potential to later fine tune these models is also a major strength compared to closed-source proprietary models. Since model size and computational efficiency are critical for large-scale screening, we compared performance across models with different parameter counts and identified the smallest model that retains strong predictive capability. We also examined whether providing cell-line–specific genetic context improves predictions by comparing a general system prompting strategy with one designed to the RPE1 *TP53* -deficient background. Furthermore, we assessed whether model performance benefits from the inclusion of external biological information, such as pathway annotations from GO databases associated with the queried genes. Finally, to test the robustness and generalizability of our findings, we validated the approach using independent Olaparib CRISPR screens conducted in multiple prostate cancer cell lines, including C4-2B, LNCaP, 22Rv1, DU145 as well as in the ovarian cancer cell line OVCAR8 [33, 34].

As a further note, we also explored the capability of the models to infer synthetic-rescue: defined as a genetic interaction where the cell is normally sensitive to a perturbation, such as drug treatment or knocked out genes independently, but when applied together, the cell becomes viable [35, 36]. We expect that this is a much harder task for an LLM compared to synthetic lethality prediction, based on previous studies [37, 38]. Therefore, we only aimed to quantify the distribution for this interaction to occur.

After we thoroughly tested the capabilities of the LLMs in this study, we constructed a list of gene pairs, where the genes have been pre-selected as clinically relevant, and exhaustively ran the best performing mid-size LLM (Qwen2.5-32B-Instruct) on a total of 398,277 pairs to obtain the LLM derived score for synthetic lethality alongside the model’s reasoning for all of them. The resulting list was further filtered by a custom LLM-assisted pipeline with internet search capability to estimate the novelty and feasibility of the predicted relationships. We made the resulting enhanced list publicly available at https://github.com/Paureel/LLMsynthlet, which we believe will complement the existing, experiment-based SL databases well [39–43]. The summary of the whole workflow can be found in Figure 1.

**Figure 1.**
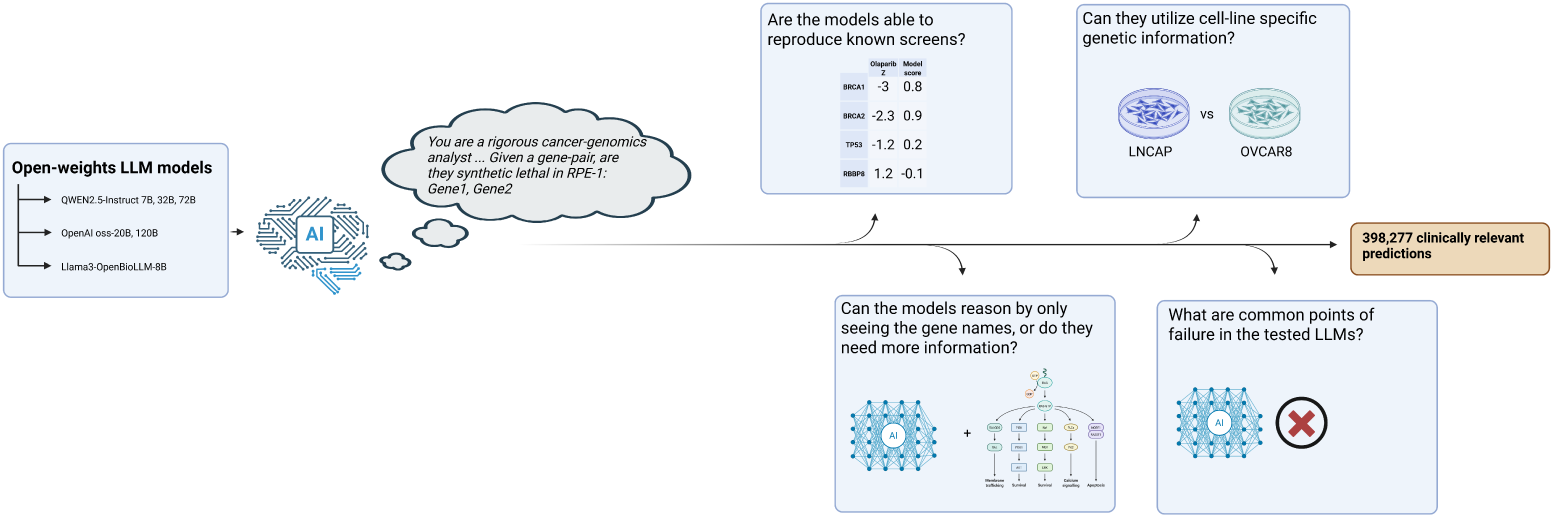
Schematic representation of the workflow in this paper. We tested multiple open-weights LLM models with a uniform prompt to reproduce known Olaparib screens. We also tested if adding information about the genes or cell lines affect the results and collected common LLM pitfalls. We also predicted 398,277 new clinically relevant pairs of genes as an explorative study.

### Pan-LLM benchmarking of CRISPR screen reproduction via zero-shot reasoning

We evaluated the ability of open-weights large language models to reproduce established genome-wide CRISPR screening results using internal knowledge and reasoning alone and then added pathway information to decide if LLMs benefit from additional gene level information.

We ran every model on all the genes present in the Olivieri M et al. dataset [3], focusing on the Olaparib treatment with additional pathway available to the models. We took the generated hypotheses alongside their assigned scores from the models, first without additional information about the genes, and then with information included. We compared the scores from the models with the ground truth information from the original screen and visualized the distribution of the scores in Figure 2A. For ROC analysis, we defined the positive class (gold standard SL hits) as the top 42 Olaparib-sensitizing genes ranked by the original normalized Z-scores, matching the authors’ reported cutoff of *z < −*3 and all remaining evaluated genes were treated as negatives. For subsequent analyses, if otherwise not stated, we used the same cutoff value. Because these benchmark sets were heavily imbalanced, we also conducted model performance evaluation using precision-recall curves with addition related metrics, shown in the supplementary information.

**Figure 2.**
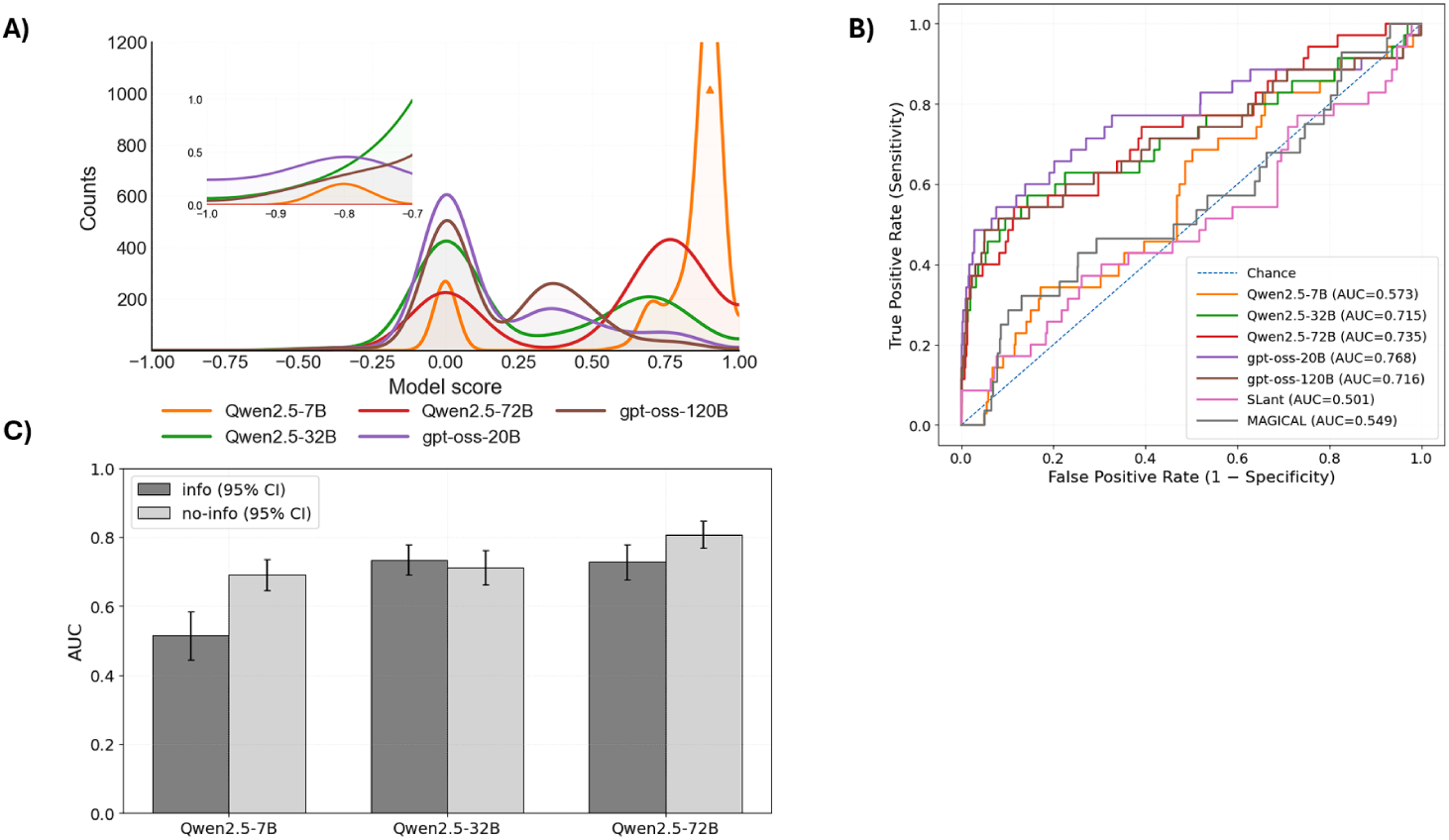
Comparative evaluation of multiple open-weights LLMs, showing predicted score distributions, ROC performance, and the effect of providing additional gene and pathway information across different model sizes. A) Distribution of the LLM predicted scores for every gene on the Olivieri M et al. dataset with additional pathway information available to the models. The inset plot highlights the distribution on the synthetic rescue end of the scores. B) ROC curves comparing the performance of different open weights LLM and non-LLM models on the Olivieri M et al. dataset. The performance of the models saturates around 0.73 AUC.C) Qwen2.5-Instruct series LLM models AUC scores across different parameter size models with and without supplying pathway information in the system prompt.

Generally, the distribution plots showed a strong bias for values for 0 and 0.8, as opposed to the expectation of having had a more uniform distribution. The Qwen2.5-7B-Instruct model predicted synthetic lethality for nearly all the genes and the Llama3-OpenBioLLM-8B model produced very inconsistent results when it came to scoring and reasoning, often just repeating words, and therefore was removed from further analysis (68.84% empty responses, 81.83% generations hitting the 8000-token cap, 91.36% failing the expected parseable output format, and 18.62% showing severe repetition). The ability of our workflow to extract the reasoning and the corresponding scores were robust, with unsuccessful extractions (gene names or score not found) occurred 0.017% for Qwen2.5-7B-Instruct, 0.006% for Qwen2.5-32B-Instruct, 0.03% for Qwen2.5-72B-Instruct, 1.15% for gpt-oss-20B and 0% for gpt-oss-120B. Apart from a random baseline, we also assessed the performance of non-LLM-based models in the comparison. We included SLant (random forest classifier trained on cross-species protein-protein interaction network topology features) and MAGICAL (multi-class random forest model built from protein-protein interaction network topology features) and applied them on the same problem and evaluation technique [44, 45].

In Figure 2B. The ROC curves of the models can be seen. Overall, most models performed better than random and non-LLM models(Table 1). The performance seemed to saturate at the 32B range, where the 7B model did not produce a usable output (mostly predicted lethal for everything), and the 72B model was not economic to run on hundreds of thousands of examples, since it could not be sharded across 8 GPU-s using data parallelism.

**Table 1.** Performance comparison of evaluated predictors on the Olivieri M et al. dataset. Area under the ROC curve (AUC), F1 score, and balanced accuracy are reported.

| model | AUC | f1 | balanced accuracy | optimal threshold |
| --- | --- | --- | --- | --- |
| gpt-oss-20B | 0.768 | 0.044 | 0.733 | 0.3 |
| Qwen2.5-72B | 0.735 | 0.012 | 0.664 | 0.8 |
| Qwen2.5-32B | 0.715 | 0.017 | 0.695 | 0.7 |
| gpt-oss-120B | 0.715 | 0.036 | 0.709 | 0.55 |
| Qwen2.5-7B | 0.573 | 0.008 | 0.571 | 0.9 |
| MAGICAL | 0.55 | 0.016 | 0.584 | 0.19 |
| Random baseline | 0.502 | 0.006 | 0.500 | 0 |
| SLant | 0.501 | 0.06 | 0.5 | 0 |

We also investigated whether providing curated pathway and genetic annotations improved LLM performance in predicting CRISPR screening outcomes. For this part we only focused on the Qwen2.5-Instruct models, since they performed as good as the gpt-oss models, but were much more efficient to run. In Figure 2C it can be observed that there was no significant difference (P = 0.25, two-tailed paired Wilcoxon signed-rank test) between pathway information enriched prompt (info) and without this addition (no-info), except in the case of the 7B model where we observed a trend on supplying pathway and genetic information degraded its performance. Interestingly, the repeated measurements on different subsets of the input gene list affected minimally the variance in performance, hence the small error bars. Since we did not observe significant difference between supplying pathway and genetic information or not on the system prompt for the cases with better than random performance, in the consecutive analyses we always used adding those additional information.

### Generalization of LLM-based synthetic lethality predictions across genome-wide CRISPR screening datasets

We tested the transferability of our method on the Tsujino T et al. paper’s dataset, involving 4 prostate cancer cell lines. This also allowed us to supply the genetic background of the cell-lines in the system prompt and see if it improves the predictions. For this part we only used the Qwen2.5-Instruct-32B model, since it was deemed the best performing given the computational efficiency of running it.

The comparison of performance across different cell lines and if the genetic information was added to the system prompt can be seen in Figure 3A, with the corresponding precision-recall curves and average precision values shown in Supplementary Figure 2.. Interestingly, our results indicated that adding genetic background information decreases the performance of the models. This effect could have been due to the fact, that the LLM can get confused by the richer input information and predicts false positives more often, as this can be deduced from the distribution plots from Figure 3B.

**Figure 3.**
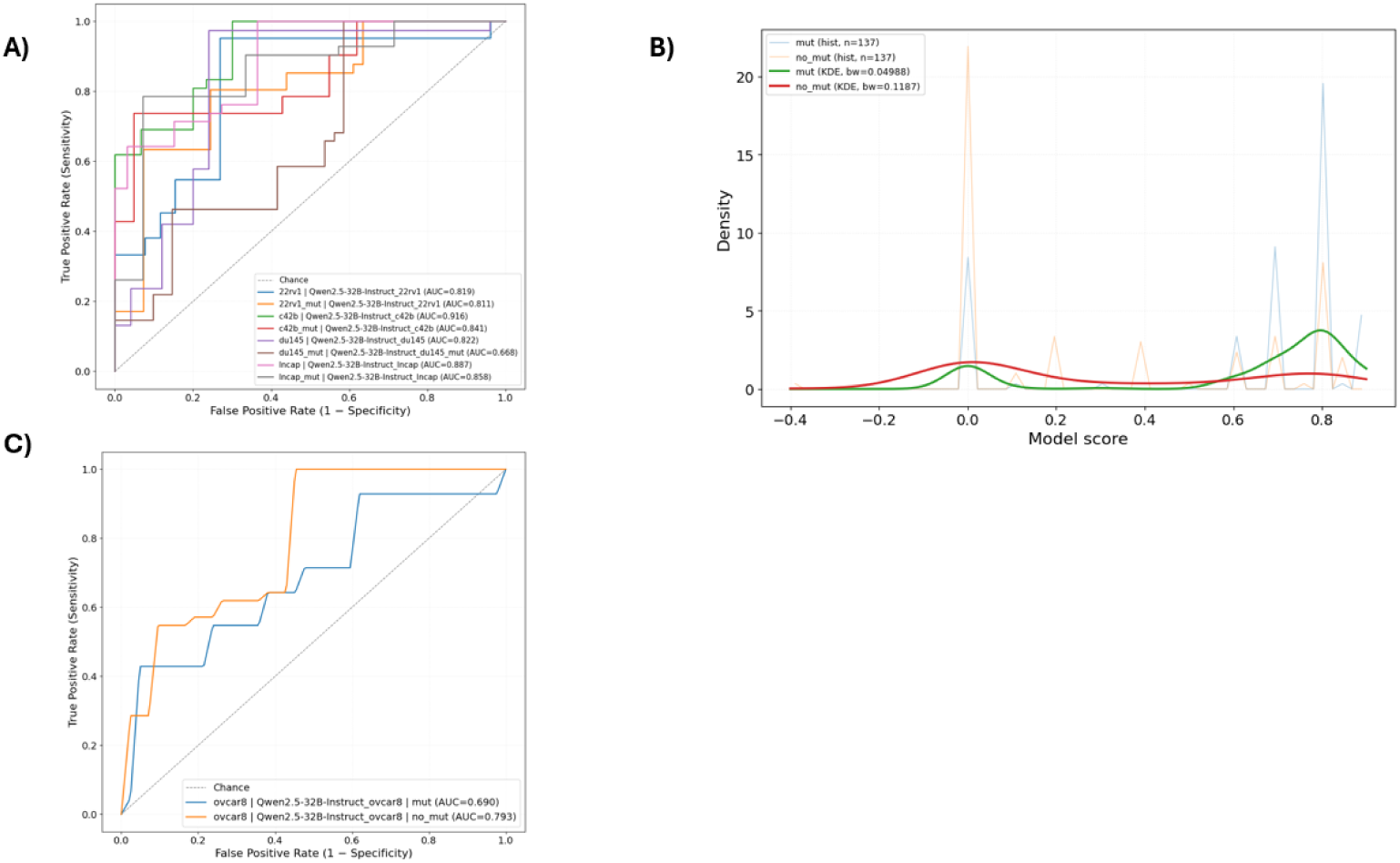
Qwen2.5-Instruct-32B models reproduced Tsujino T et al. and Liu et al. paper’s dataset. A) The ROC curves were compared across cell-lines and if the model’s system prompt contained genetic information or not for the Tsujino T et al. dataset. B) Distribution of model output score for the 22RV1 cell line as a representative example, showing the difference between with or without additional genetic information added to the system prompt. C) ROC curves across different prompting techniques for the Liu et al. paper’s dataset.

We performed similar comparison in the Liu et al. paper’s dataset. The main advantage of this comparison was that it allowed us to test our approach on a dataset that was released after the tested model’s cut-off data, eliminating data leakage entirely. Our results on this dataset indicated that both the mut and no-mut version of the prompting resulted in better than random prediction, but altogether lower scores compared to the Tsujino T et al. paper’s dataset (Figure 3C, precision-recall analysis for OVCAR8 is shown in Supplementary Figure 3).

### Distribution and filtering of clinically relevant two-gene synthetic lethality predictions

After performing the analyses of the predictability of the models, we characterized the global score distribution of all putative clinically relevant pairwise SL interactions predicted for the RPE-1 cell line and devised a simple LLM workflow to further filter the candidate SL relationships (Figure 4A).

**Figure 4.**
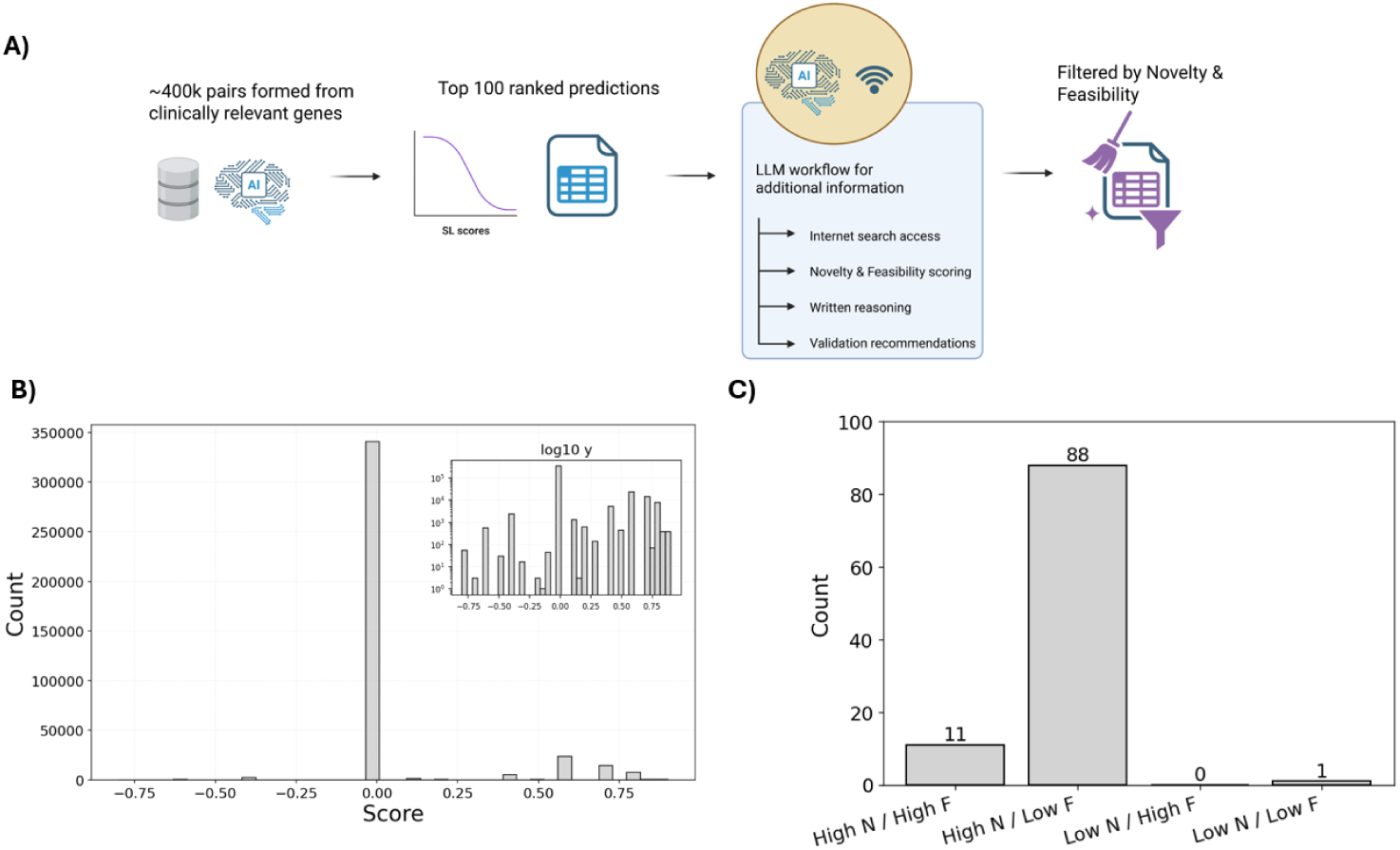
Overview of the post-prediction prioritization pipeline and its outcomes. A) Schematic representation for scoring gene–gene pairs, selecting top hits, and using an LLM-assisted workflow to assign novelty and feasibility and to recommend validations. B) LLM Score distribution across all 398,277 predictions, with most predictions centered at 0 and a smaller high-scoring SL tail (log-scale inset). C) Counts of the top 100 predicted SL pairs binned by LLM-assigned novelty (N) and feasibility (F) categories.

Most importantly, the predictions concentrated on exactly the zero value (more than 80%), meaning that most gene-gene pairs have been predicted as non-synthetic lethal. The model produced a smaller subset of gene pairs centered around 0.55 predicted as SL (less than 20%), that can be used for further prioritizing and filtering. Negative score values are rare, but they do exist, meaning the model has predicted a small subset of synthetic rescue relationships (less than 2%) (Figure 4B). The shape of the distribution implied that the results are better used downstream for ranking, prioritizing the extreme tails, and not for broad classification.

Next, we ordered the predictions by their respective LLM assigned score and filtered for the top 100 most synthetically lethal predicted gene pairs. We used a custom-made LLM-assisted workflow (detailed in Materials and Methods) to gather additional information on the given gene-pair to estimate the novelty and feasibility of the given interaction, alongside a written reasoning and validation recommendations. The LLM scored the novelty and feasibility by assigning a score from 0 to 10 for both, where 0 means a known interaction for novelty and a biologically and/or experimentally impossible relationship. We quantified the results by further categorizing them to 4 categories of high novelty – high feasibility, low novelty – high feasibility, high novelty – low feasibility and low novelty – low feasibility (cut-off value at 5 for both). The resulting categorization indicated that judged by the LLM, only 11 of predicted interactions ended up in high novelty – high feasibility bin, while most were predicted as high novelty but low feasibility (Figure 4C).

Interestingly, in the top 100 predictions the *TBP* (TATA-box binding protein) gene was vastly overrepresented (73/100). While technically correct, the enrichment of this gene in the final result is likely attributed to *TBP* being biologically central, hub-like component, as it is being part of the transcription initiation machinery and is described as essential at virtually every eukaryotic promoter [46]. This makes the predictions related to *TBP* biologically plausible, but non-selective.

To investigate the possibility whether the LLM acts as a detector for genes that are lethal even when knocked out separately, apart from their counterpart in the proposed synthetic lethal pair, we compared all predictions for the 398,277 gene pairs against mean RPE-1 single-gene essentiality from the DepMap CRISPR gene-effect data (Supplementary Figure 4). The results showed that the association for the higher gene-essentiality from the pair and the calculated synthetic lethality score from the LLM was 0.129 (Spearman correlation). This suggests that single-gene essentiality contributes, but only in a modest way, without explaining the ranking on its own.

### Observed error modes in LLM outputs

We also collected some common LLM reasoning and instruction-following errors. One frequently (around 10%) appearing issue in the cell line genetic background setting was that the LLM in its output tried to override the system prompt. For example, after defining the input interaction as Olaparib – *FANCM*, it outputted: `I will assume a hypothetical gene interaction with BRCA2"` instead of Olaparib treatment or *PARP1*.

Another problem occurred when the LLM mixed up the functions of genes, that are named similarly to each other. For example, in one case the LLM reasoned that *CtIP* is ubiquitylated by *KLHL18*, when according to the latest research, it is ubiquitylated by *CUL3* -*KLHL15* [47, 48]. We suspect that this problem can be traced back to how the gene names are tokenized, but it remains unconfirmed.

Lastly, we noticed that the LLM, when making predictions using background genetic information given a specific cell line, often determined that the cell has so many HR-related genes deleted already, that it automatically assigned synthetic lethality without further reasoning. This significantly elevated the false positive predictions and rendered the ability of the LLM incorporating conditionals more difficult.

## Discussion

Our goal in this study was to test whether popular open-weights LLMs can reproduce known SL interactions from publicly available genome-wide CRISPR treatment screens, using only the internal biological knowledge of the models. Comparing different models and prompting strategies, the best performing models successfully and reliably reproduced several Olaparib CRISPR screen hits across several cell lines better than random guessing, indicating that LLMs could be used as practical filters for large in silico screens. This way they might be used to prioritize a manageable subset of gene pairs from millions of possibilities. We also stress that the evaluation of the models has been done in a zero-shot manner with a small formatting few-shot scaffold included in the system prompt. The models were not fine-tuned or parameter tuned, and only prompt engineering was used to make the LLMs output the required structure. For this we added three specific examples of synthetic lethal, neutral and synthetic rescue examples, that were not used in further evaluations, meaning we did not evaluate on these gene pairs, and they were only used as examples to help align the LLM’s output. In this study, we also included both “reasoning” models such as gpt-oss-20b, gpt-oss-120b (reasoning effort set to medium) and “non-reasoning” models, which included Llama3-OpenBioLLM-8B, Qwen2.5-7B-Instruct, Qwen2.5-32B-Instruct, Qwen2.5-72B-Instruct. However, the difference between the performance of these two model types was small when we compared the models with similar parameter size, and it is unclear how the built-in reasoning mode helps with the final reasoning output of the models.

During our analysis, several limitations became clear. First, our selection of “ground truth” by taking the cut-off values from the original papers after ordering the genes by sensitizing z-values can be somewhat arbitrary and includes several evident false positives for specific cell lines (*IL22* and *ESR1* in RPE1 for example [49]) due to off-target effects [50–52]. However, most tested models classified these as neutral and not SL relationship, correctly. This suggests that LLM-based evaluation and reasoning may help flag incorrect hits. Second, Olaparib targets several other proteins, and is not just a pure PARP1 inhibitor, and our prompting focus on *PARP1* alone might not fully capture the full target and pathway effects in the cells [53–55]. We observed a strong link between parameter size of the model and their performance, where fewer parameter models performed and followed instructions worse, often collapsing the score distribution, and this relationship quickly saturated as a function of increasing parameter size. Modifying the prompt by adding pathway and gene specific information as well as adding information on the genetic background of the tested cell lines contributed little to the predictive behavior of the models. This suggests that current LLMs already learn enough information on their training phase to make adequate predictions and more structural integration of external resources might be needed to further boost their performance.

We observed that the predicted score distributions by the LLMs are often highly binned, often centered around values like 0 and 0.75 as opposed to our expectation of a smooth distribution between –1 and 1. We suspect that this effect comes from the way we defined the scoring method in the system prompt, and generally should not affect the predictive power of the models, since we used the ranking of the genes to evaluate the results. Interestingly, we did observe negative scores indicative of synthetic rescue relationships, pointing toward capabilities in the model to indicate resistance mechanisms to PARP inhibition that are difficult to discover experimentally. Overall, this work should be viewed as exploratory and the first of its kind to use LLMs on high throughput in silico synthetic lethality prediction. We argue that the enrichment of known SL pairs across three independent studies, the identification of likely false positives in the datasets and the capability to score nearly 400000 clinically relevant gene pairs in RPE-1 cell line point into the direction of LLM-based reasoning can complement existing graph- and expression-based methods. It should be noted that we do not resolve to what extent the tested LLMs are using genuine biological reasoning as opposed to preferential treatment of well-annotated genes, and further validation needs to be conducted to resolve this. It is also important to note that the LLM-assisted prioritization of the predicted clinically relevant gene pairs is a highly explorative step which requires further benchmarking. Importantly, the LLM-derived predictions in the final set were not validated experimentally, in this study and therefore should be interpreted as hypotheses that required further wet-lab confirmation.

During downstream analysis we noticed that a significant subset of issues arise due to the incomplete instruction following of the LLMs, highlighting the known fact that these models are sensitive to variations in the prompts [56–58]. Further studies should include rigid testing of different prompts to quantify this sensitivity.

We are also aware of the possibility of potential data leakage, since some of the papers have been published before the cut-off date of the LLM’s pre-training phase, similarly to other fields [59, 60]. For this reason, we have included the Liu et al. paper, that has been made available after the cut-off date. We also measured a high false positive rate across all models, that might be an indication of the models not memorizing the data. Finally, we compared tens of thousands of genes with measured values from the papers, and it is highly unlikely that the actual spreadsheet with the values have been included in the pre-training run for these models. Nevertheless, we reserve the possibility of minimal data contamination through the means of the papers we have tested the LLMs on being in their pre-training dataset, or through the LLM prioritizing overrepresented genes from the same place. Another source of data leakage could have come from the inference runs where we included GO annotations. However, GO augmentation had minimal impact on benchmark performance in our ablation tests, suggesting that this had minimal impact on the results.

Future directions include the expansion of the analysis for further treatments and expanding the system prompt to accept combinatorically higher combinations of genes (*≥* 3 genes), further analyzing synthetic rescue predictions and tool calling to combine LLM reasoning with other modalities, such as structural biology databases in a more principled way. Another interesting future direction would be to conduct a similar analysis, but via state-of-the-art, proprietary LLMs through API-s and directly compare their capabilities to open-weights models.

## Materials and Methods

We defined as and regarding to synthetic lethality in the paper for ease as both the interaction between genes if both are deleted (functional SL) and deleted gene + drug treatment (chemogenetic interaction), and in the same way for synthetic rescue. In the latter case we specified the main target of the treatment as a gene.

We acquired data to reproduce and validate our approach from the Olivieri M et al. [3], Tsujino T et al. [34] and Liu et al. [33] papers. The exact papers are listed in Table 2. In Olivieri M et al., 31 genome-scale CRISPR/Cas9 dropout screens were performed in *TP53* -deficient RPE-1 cells across 27 genotoxic compounds. The authors used normalized Z-scores to order the genes regarding their sensitization potential. In Tsujino T et al., the authors performed similarly a genome-scale CRISPR/Cas9 dropout screen in four *BRCA1/2* proficient prostate cancer cell lines (LNCAP, C42B, 22RV1, DU145). The cell lines were treated with Olaparib versus DMSO, then the genes have been ordered by sgRNA depletion scores. In Liu et al. the authors performed a genome-wide CRISPR/Cas9 knockout dropout screen in the *BRCA1/2* –wild-type ovarian cancer cell line OVCAR8, comparing 2-week Olaparib treatment versus DMSO with Day 0 as baseline and ranking genes by sgRNA depletion using MAGeCK/MAGeCK-VISPR. We took the data from these experiments, and our processing steps included filtering and ordering the data by Olaparib sensitization in both cases using the scores for ordering the genes from the original experiments. When constructing the benchmarks, we used the author-defined positive and negative sets for Olaparib-gene knockout list. Where this information was not supplied, we used the default definition from Olivieri M et al. taking the top 42 genes as positive, and the rest of the genes as negatives. This resulted in a comparison with different number of ground truth samples, and the calculation of the metrics were conducted independently on each dataset’s own values. The exact number were the followings: 42 positives and 98 negatives for 22Rv1, 42 positives and 168 negatives for C4-2B, 38 positives and 142 negatives for DU145, 42 positives and 155 negatives for LNCaP, 42 positives and 42 negatives for OVCAR8, and 42 positives and 11,610 negatives for the Olivieri M et al dataset.

**Table 2.**
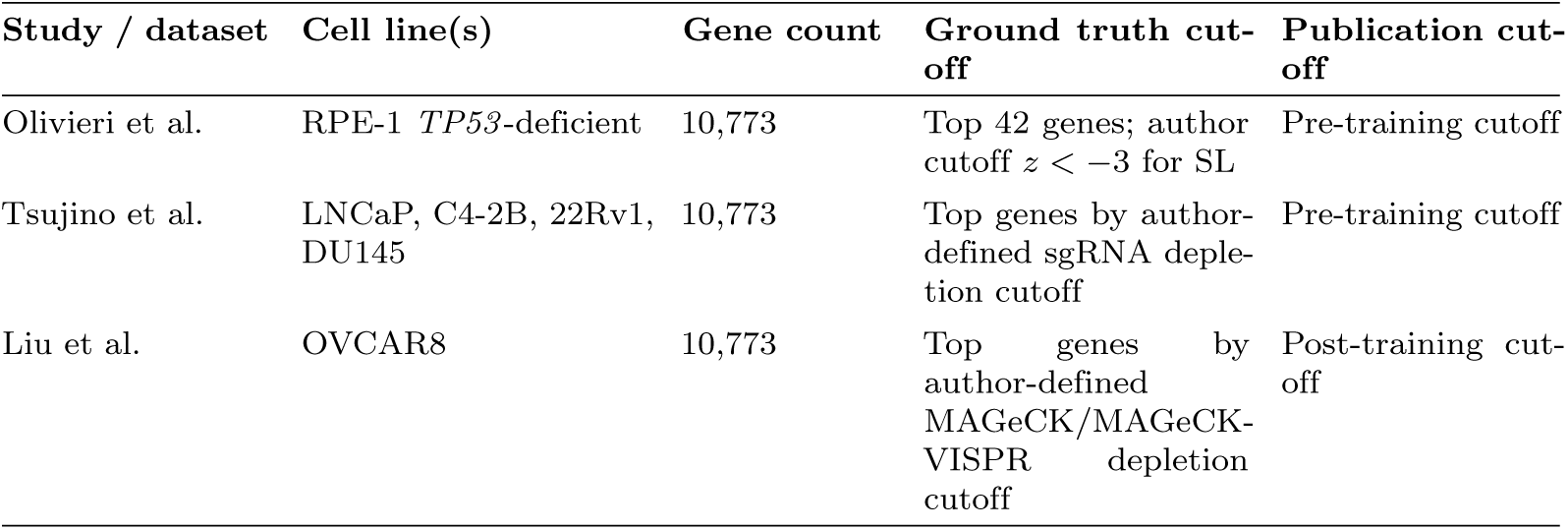
Summary of the CRISPR screens used in this study. Gene count refers to the number of genes evaluated in our downstream analysis. For all datasets, after filtering, altogether 10,773 genes were evaluated after preprocessing and filtering. The Liu et al. dataset was published after the training cutoff of the tested models and was therefore used as a post-training-cutoff benchmark.

For the in silico experiments we constructed the prompts the LLMs were iterating on by defining a system prompt, which was then used on all gene pairs or treatment-single genes. For further information refer to the exact prompts in the supplementary information. The system prompt was appended in front of the gene-pair or treatment-single gene specific prompts, which followed the format: `PARP1 (Olaparib treatment), Gene1 { additional information on Gene1`. For the prospective screen we used the format: `Gene1, Gene2 { additional information on Gene1, additional information on Gene2`. The system prompts could also contain cell-line specific information on the exact genetic background of the cell lines, which was obtained by adding *TP53* mutation in RPE-1 cell line case and was obtained in the following way for the 22RV1, LNCAP, C42B and DU145 cell lines: we filtered for genes indicated in DNA repair and included SNP-s with HIGH VEP IMPACT and included INDELS with HIGH VEP IMPACT. The resulting gene list got listed in the system prompts for the corresponding cell lines. In the data the Liu et al. paper is based on (OVCAR8), the OmicsSomaticMutationsMatrixDamaging.csv file from DepMap (version 25Q3) [15] was used to filter for damaging mutations only, and the resulting gene list was added to the system prompt. All statistical tests were conducted with alpha = 0.05.

To supplement additional information to the LLMs, the Gene Ontology database go.obo file was joined with the Gene Ontology Annotation goa human.gaf file [61, 62]. We included all available annotations without further filtering. This allowed us to add richer information about the genes regarding their interactions and pathways they are part of, helping the LLM to reason. We used the Gene Ontology database, as opposed to other popular databases such as Reactome and KEGG, because it provided a standardized, machine-readable summaries for each gene’s functions and localization used in the study, that fit the per-gene prompt design. The output of the LLMs was capped to 8000 output tokens, and this way any additional information not fitting to this window was truncated.

Given the same definition for resistance (¿6), there have not been any genes contributing to resistance in this dataset after checking the distribution. We also removed genes that were below 40th percentile expressed using TPM values from DepMap (version 25Q3) RPE-1 cell lines, since these could contribute to false positives (CRISPR off-target effects [50–52]) in the original measurements if ranked high as sensitizing genes. This cutoff was determined as a conservative low-expression filter that still allowed us to retain most of the genes, and is based on a representative dropout screen analysis [63]. We then evaluated all the 10773 genes as `PARP1 (Olaparib sensitivity) - Gene X` in a pairwise manner. The LLM was tasked to summarize the genes, give a mechanistic interpretation of the possible synthetic lethality and assign score from –1 to 1 where –1 is strong synthetic rescue, 0 is neutral/addition effect and 1 is strong synthetic lethality. The output from the LLM was a continuous text, and the scores were extracted from it by pattern matching. Since the LLM score output is not calibrated, we also did this step also to calibrate the scoring. For determining the optimal cutoffs we used Youden’s J. For the comparison to a random baseline, we randomly permuted each model’s score output on each gene, which allowed us to preserve the same distribution, while breaking any association with the ground truth data.

For comparing the LLM models to see if the supplied pathway information improves the results, we only sampled 2000 genes from the Olivieri M et al. screen in a way, that by ordering the genes by z-values for Olaparib, we selected the top 1000 sensitizing genes, and we randomly sampled 1000 more from the rest of the genes, ensuring that the models can be run efficiently on multiple replicates, from which we used 3 replicates per model for this part of the analysis. Since by using three replicates with the sampling described previously yielded in small between-replicate variance, we constructed the error bars representing the 95% CIs on the group mean using a stratified bootstrap over genes within each replicate by resampling positives/negatives with replacement to estimate within-run variance and then combined with between-run variance via variance propagation.

The LLMs used in this study were run with their recommended hyperparameters given their model cards. The gpt-oss-20b and gpt-oss-120b models were ran on medium reasoning mode. We used the Hugging Face Transformers library for running the LLMs (version 4.56.2) [64]. All inference runs were performed on one DGX node consisting of 8x NVIDIA H100 SXM (80 GB) GPUs on Gefion, Denmark’s national AI supercomputer [65]. We used data parallel inference on models that fit into one GPU, and for the gpt-oss-120b model we used intra-node model parallelism. For building the LLM-assisted pipeline to further filter the top 100 SL predictions we used the eu.anthropic.claude-sonnet-4-20250514-v1:0 model. The pipeline consisted of an iterative search on dynamic keywords with Tavily (version 0.7.19) with maximum retrieved results set to 20 and an LLM interpretation part. The exact system prompt can be found in the supplementary information alongside the exact keywords used.

The non-LLM baseline models have been obtained from their respective repositories and were run with their default parameters [44, 45]. In the case of SLant, we included the pairs of genes predicted as synthetic lethal that were paired with *PARP1*. For MAGICAL, we first ordered the results for all pairwise *PARP1* predictions by SL score from the model and used the resulting ordered list to compare against the ground truths.

## Data Availability

Benchmark input data were obtained from the cited published studies and public resources named in Materials and Methods. The results from the run involving the 398,277 gene pairs are available at https://github.com/Paureel/LLMsynthlet.

## Code Availability

The code used for the inference of the LLM models can be accessed at https://github.com/Paureel/LLMsynthlet.

## Acknowledgments

Figures 1 and 4A were created with BioRender.com.

The project was funded by Novo Nordisk Foundation (NNF25OC0104818 to Z.S.), Breast Cancer Research Foundation (BCRF-23-159 to Z.S.), NIH Grant 1 P01 CA228696-01A1 to Z.S, Kræftens Bekæmpelse (R325-A18809 and R342-A19788 to Z.S.), Det Frie Forskningsråd Sundhed og Sygdom (2034-00205B to Z.S.), Basser Foundation (to Z.S.), Susan G Komen Foundation (to Z.S.). This work was also supported by the Ovarian Cancer Research Alliance by grant CRDGAI-2025-3-1992 to Z.S. and I.C., and National Research, Development, and Innovation Office of Hungary (NKKP-153428) to I.C.. The funders played no role in study design, data collection, analysis, interpretation of data, or writing of the manuscript.

## Author Contributions

Conceptualization, A.P., Z.Sz., M.D., B.Z., I.C., Z.S, O.K.; Methodology, A.P., B.Z., I.C., Z.S.; Software A.P.; Formal analysis A.P., Writing A.P., Z.S.; Funding acquisition I.C., Z.S..

## References

1. Nijman SMB. Synthetic lethality: General principles, utility and detection using genetic screens in human cells. FEBS Letters. 2011 Jan;585(1):1–6. Available from: http://doi.wiley.com/10.1016/j.febslet.2010.11.024.

2. Lord CJ, Ashworth A. PARP inhibitors: Synthetic lethality in the clinic. Science. 2017 Mar;355(6330):1152–8. Available from: https://www.science.org/doi/10.1126/science.aam7344.

3. Olivieri M, Cho T, Álvarez Quilón A, Li K, Schellenberg MJ, Zimmermann M, et al. A Genetic Map of the Response to DNA Damage in Human Cells. Cell. 2020 Jul;182(2):481–96.e21. Available from: https://linkinghub.elsevier.com/retrieve/pii/S0092867420306735.

4. Zimmermann M, Murina O, Reijns MAM, Agathanggelou A, Challis R, Tarnauskaite z, et al. CRISPR screens identify genomic ribonucleotides as a source of PARP-trapping lesions. Nature. 2018 Jul;559(7713):285–9. Available from: https://www.nature.com/articles/s41586-018-0291-z.

5. Ryan CJ, Devakumar LPS, Pettitt SJ, Lord CJ. Complex synthetic lethality in cancer. Nature Genetics. 2023 Dec;55(12):2039–48. Available from: https://www.nature.com/articles/s41588-023-01557-x.

6. Romero R, Sayin VI, Davidson SM, Bauer MR, Singh SX, LeBoeuf SE, et al. Keap1 loss promotes Kras-driven lung cancer and results in dependence on glutaminolysis. Nature Medicine. 2017 Nov;23(11):1362–8. Available from: https://www.nature.com/articles/nm.4407.

7. Burgold T, Karakoc E, Gonçalves E, Barrio-Hernandez I, Dwane L, Silva R, et al. A next-generation dual guide CRISPR system for genetic interaction library screening. Nature Communications. 2025 Dec;17(1):561. Available from: https://www.nature.com/articles/s41467-025-67256-9.

8. Zhou P, Chan BKC, Wan YK, Yuen CTL, Choi GCG, Li X, et al. A Three-Way Combinatorial CRISPR Screen for Analyzing Interactions among Druggable Targets. Cell Reports. 2020 Aug;32(6):108020. Available from: https://linkinghub.elsevier.com/retrieve/pii/S2211124720310056.

9. Manshadi MD, Setoodeh P, Ramezani A, Rajabzadeh AR, Zare H. Higher order synthetic lethals are keys to minimize cancer treatment effects on non-tumor cells. Systems Biology; 2025. Available from: http://biorxiv.org/lookup/doi/10.1101/2025.01.31.635848.

10. Schäffer AA, Chung Y, Kammula AV, Ruppin E, Lee JS. A systematic analysis of the landscape of synthetic lethality-driven precision oncology. Med. 2024 Jan;5(1):73–89.e9. Available from: https://linkinghub.elsevier.com/retrieve/pii/S2666634023004075.

11. Shen JP, Ideker T. Synthetic Lethal Networks for Precision Oncology: Promises and Pitfalls. Journal of Molecular Biology. 2018 Sep;430(18):2900–12. Available from: https://linkinghub.elsevier.com/retrieve/pii/S0022283618306338.

12. Ngoi NYL, Gallo D, Torrado C, Nardo M, Durocher D, Yap TA. Synthetic lethal strategies for the development of cancer therapeutics. Nature Reviews Clinical Oncology. 2025 Jan;22(1):46–64. Available from: https://www.nature.com/articles/s41571-024-00966-z.

13. Henkel L, Rauscher B, Boutros M. Context-dependent genetic interactions in cancer. Current Opinion in Genetics & Development. 2019 Feb;54:73–82. Available from: https://linkinghub.elsevier.com/retrieve/pii/S0959437X19300152.

14. Rossiter NJ, Huggler KS, Adelmann CH, Keys HR, Soens RW, Sabatini DM, et al. CRISPR screens in physiologic medium reveal conditionally essential genes in human cells. Cell Metabolism. 2021 Jun;33(6):1248–63.e9. Available from: https://linkinghub.elsevier.com/retrieve/pii/S1550413121000619.

15. Tsherniak A, Vazquez F, Montgomery PG, Weir BA, Kryukov G, Cowley GS, et al. Defining a Cancer Dependency Map. Cell. 2017 Jul;170(3):564–76.e16. Available from: https://linkinghub.elsevier.com/retrieve/pii/S0092867417306517.

16. Feng Y, Long Y, Wang H, Ouyang Y, Li Q, Wu M, et al. Benchmarking machine learning methods for synthetic lethality prediction in cancer. Nature Communications. 2024 Oct;15(1):9058. Available from: https://www.nature.com/articles/s41467-024-52900-7.

17. Huang Y, Yuan R, Li Y, Xing Z, Li J. Struct2SL: Synthetic lethality prediction based on AlphaFold2 structure information and Multilayer Perceptron. Computational and Structural Biotechnology Journal. 2025;27:1570–7. Available from: https://linkinghub.elsevier.com/retrieve/pii/S2001037025001345.

18. Zhang K, Wu M, Liu Y, Feng Y, Zheng J. KR4SL: knowledge graph reasoning for explainable prediction of synthetic lethality. Bioinformatics. 2023 Jun;39(Supplement 1):i158–67. Available from: https://academic.oup.com/bioinformatics/article/39/Supplement_1/i158/7210467.

19. Fan K, Gökbağ B, Tang S, Li S, Huang Y, Wang L, et al. Synthetic lethal connectivity and graph transformer improve synthetic lethality prediction. Briefings in Bioinformatics. 2024 Jul;25(5):bbae425. Available from: https://academic.oup.com/bib/article/doi/10.1093/bib/bbae425/7745393.

20. Seale C, Tepeli Y, Gonçalves JP. Overcoming selection bias in synthetic lethality prediction. Bioinformatics. 2022 Sep;38(18):4360–8. Available from: https://academic.oup.com/bioinformatics/article/38/18/4360/6649727.

21. Luo R, Sun L, Xia Y, Qin T, Zhang S, Poon H, et al. BioGPT: generative pre-trained transformer for biomedical text generation and mining. Briefings in Bioinformatics. 2022 Nov;23(6):bbac409. Available from: https://academic.oup.com/bib/article/doi/10.1093/bib/bbac409/6713511.

22. Lin A, Ye J, Qi C, Zhu L, Mou W, Gan W, et al. Bridging artificial intelligence and biological sciences: a comprehensive review of large language models in bioinformatics. Briefings in Bioinformatics. 2025 Jul;26(4):bbaf357. Available from: https://academic.oup.com/bib/article/doi/10.1093/bib/bbaf357/8212018.

23. Chen Y, Zou J. GenePT: A Simple But Effective Foundation Model for Genes and Cells Built From ChatGPT. Bioinformatics; 2023. Available from: http://biorxiv.org/lookup/doi/10.1101/2023.10.16.562533.

24. Bryan JG, Niu H, Li D. Incorporating Large Language Model-Derived Information into Hypothesis Testing for Genomics. Bioinformatics; 2025. Available from: http://biorxiv.org/lookup/doi/10.1101/2025.04.30.651464.

25. Shang X, Liao X, Ji Z, Hou W. Benchmarking large language models for genomic knowledge with GeneTuring. Briefings in Bioinformatics. 2025 Aug;26(5):bbaf492. Available from: https://academic.oup.com/bib/article/doi/10.1093/bib/bbaf492/8261762.

26. Schaefer M, Peneder P, Malzl D, Lombardo SD, Peycheva M, Burton J, et al. Multimodal learning enables chat-based exploration of single-cell data. Nature Biotechnology. 2025 Nov. Available from: https://www.nature.com/articles/s41587-025-02857-9.

27. Jin Q, Yang Y, Chen Q, Lu Z. GeneGPT: augmenting large language models with domain tools for improved access to biomedical information. Bioinformatics. 2024 Feb;40(2):btae075. Available from: https://academic.oup.com/bioinformatics/article/doi/10.1093/bioinformatics/btae075/7606338.

28. Wang Z, Jin Q, Wei CH, Tian S, Lai PT, Zhu Q, et al. GeneAgent: self-verification language agent for gene-set analysis using domain databases. Nature Methods. 2025 Aug;22(8):1677–85. Available from: https://www.nature.com/articles/s41592-025-02748-6.

29. Feng Y, Zhou L, Ma C, Zheng Y, He R, Li Y. Knowledge graph–based thought: a knowledge graph–enhanced LLM framework for pan-cancer question answering. GigaScience. 2025 Jan;14:giae082. Available from: https://academic.oup.com/gigascience/article/doi/10.1093/gigascience/giae082/7943459.

30. Pal A, Sankarasubbu M. OpenBioLLMs: Advancing Open-Source Large Language Models for Healthcare and Life Sciences; 2024. Available from: https://huggingface.co/aaditya/OpenBioLLM-Llama3-70B.

31. Qwen, Yang A, Yang B, Zhang B, Hui B, Zheng B, et al.. Qwen2.5 Technical Report. arXiv; 2024. Version Number: 2. Available from: https://arxiv.org/abs/2412.15115.

32. OpenAI, Agarwal S, Ahmad L, Ai J, Altman S, Applebaum A, et al.. gpt-oss-120b & gpt-oss-20b Model Card. arXiv; 2025. Version Number: 1. Available from: https://arxiv.org/abs/2508.10925.

33. Liu C, Xu F, Wu Y, Li J, Ni M, Xia S, et al. Genome-wide CRISPR-Cas9 screening identifies CLK1 inhibition as a strategy to restore PARP inhibitor sensitivity via ERCC1 isoform switching. Protein & Cell. 2025 Nov:pwaf091. Available from: https://academic.oup.com/proteincell/advance-article/doi/10.1093/procel/pwaf091/8314228.

34. Tsujino T, Takai T, Hinohara K, Gui F, Tsutsumi T, Bai X, et al. CRISPR screens reveal genetic determinants of PARP inhibitor sensitivity and resistance in prostate cancer. Nature Communications. 2023 Jan;14(1):252. Available from: https://www.nature.com/articles/s41467-023-35880-y.

35. Liu M, Dong Q, Chen B, Liu K, Zhao Z, Wang Y, et al. Synthetic viability induces resistance to immune checkpoint inhibitors in cancer cells. British Journal of Cancer. 2023 Oct;129(8):1339–49. Available from: https://www.nature.com/articles/s41416-023-02404-w.

36. Bertlin JAC, Pauzaite T, Liang Q, Wit N, Williamson JC, Sia JJ, et al.. VHL synthetic lethality screens uncover CBF-b as a negative regulator of STING. Cancer Biology; 2024. Available from: http://biorxiv.org/lookup/doi/10.1101/2024.09.03.610968.

37. Sahu AD, S Lee J, Wang Z, Zhang G, Iglesias-Bartolome R, Tian T, et al. Genome-wide prediction of synthetic rescue mediators of resistance to targeted and immunotherapy. Molecular Systems Biology. 2019 Mar;15(3):e8323. Available from: https://link.springer.com/article/10.15252/msb.20188323.

38. Shen JP, Zhao D, Sasik R, Luebeck J, Birmingham A, Bojorquez-Gomez A, et al. Combinatorial CRISPR–Cas9 screens for de novo mapping of genetic interactions. Nature Methods. 2017 Jun;14(6):573–6. Available from: https://www.nature.com/articles/nmeth.4225.

39. Wang J, Zhang Q, Han J, Zhao Y, Zhao C, Yan B, et al. Computational methods, databases and tools for synthetic lethality prediction. Briefings in Bioinformatics. 2022 May;23(3):bbac106. Available from: https://academic.oup.com/bib/article/doi/10.1093/bib/bbac106/6555403.

40. Guo J, Liu H, Zheng J. SynLethDB: synthetic lethality database toward discovery of selective and sensitive anticancer drug targets. Nucleic Acids Research. 2016 Jan;44(D1):D1011–7. Available from: https://academic.oup.com/nar/article-lookup/doi/10.1093/nar/gkv1108.

41. Guo L, Dou Y, Xia D, Yin Z, Xiang Y, Luo L, et al. SLOAD: a comprehensive database of cancer-specific synthetic lethal interactions for precision cancer therapy via multi-omics analysis. Database. 2022 Aug;2022:baac075. Available from: https://academic.oup.com/database/article/doi/10.1093/database/baac075/6677988.

42. Zhang B, Tang C, Yao Y, Chen X, Zhou C, Wei Z, et al. The tumor therapy landscape of synthetic lethality. Nature Communications. 2021 Feb;12(1):1275. Available from: https://www.nature.com/articles/s41467-021-21544-2.

43. Li Xj, Mishra SK, Wu M, Zhang F, Zheng J. Syn-Lethality: An Integrative Knowledge Base of Synthetic Lethality towards Discovery of Selective Anticancer Therapies. BioMed Research International. 2014;2014:1–7. Available from: http://www.hindawi.com/journals/bmri/2014/196034/.

44. Dey A, Mudunuri S, Kiran M. MAGICAL: A multi-class classifier to predict synthetic lethal and viable interactions using protein-protein interaction network. PLOS Computational Biology. 2024 Aug;20(8):e1012336. Available from: https://dx.plos.org/10.1371/journal.pcbi.1012336.

45. Benstead-Hume G, Chen X, Hopkins SR, Lane KA, Downs JA, Pearl FMG. Predicting synthetic lethal interactions using conserved patterns in protein interaction networks. PLOS Computational Biology. 2019 Apr;15(4):e1006888. Available from: https://dx.plos.org/10.1371/journal.pcbi.1006888.

46. Akhtar W, Veenstra GJC. TBP-related factors: a paradigm of diversity in transcription initiation. Cell & Bioscience. 2011;1(1):23. Available from: http://cellandbioscience.biomedcentral.com/articles/10.1186/ 2045-3701-1-23.

47. Ferretti LP, Himmels SF, Trenner A, Walker C, Von Aesch C, Eggenschwiler A, et al. Cullin3-KLHL15 ubiquitin ligase mediates CtIP protein turnover to fine-tune DNA-end resection. Nature Communications. 2016 Aug;7(1):12628. Available from: https://www.nature.com/articles/ncomms12628.

48. Moghe S, Jiang F, Miura Y, Cerny RL, Tsai MY, Furukawa M. The CUL3-KLHL18 ligase regulates mitotic entry and ubiquitylates Aurora-A. Biology Open. 2012 Feb;1(2):82–91. Available from: https://journals.biologists.com/bio/article/1/2/82/818/The-CUL3-KLHL18-ligase-regulates-mitotic-entry-and.

49. Perusina Lanfranca M, Lin Y, Fang J, Zou W, Frankel T. Biological and pathological activities of interleukin-22. Journal of Molecular Medicine. 2016 May;94(5):523–34. Available from: http://link.springer.com/10.1007/s00109-016-1391-6.

50. Sheel A, Xue W. Genomic Amplifications Cause False Positives in CRISPR Screens. Cancer Discovery. 2016 Aug;6(8):824–6. Available from: https://aacrjournals.org/cancerdiscovery/article/6/8/824/5788/Genomic-Amplifications-Cause-False-Positives-in.

51. DeWeirdt PC, Sangree AK, Hanna RE, Sanson KR, Hegde M, Strand C, et al. Genetic screens in isogenic mammalian cell lines without single cell cloning. Nature Communications. 2020 Feb;11(1):752. Available from: https://www.nature.com/articles/s41467-020-14620-6.

52. Perez AR, Sala L, Perez RK, Vidigal JA. Computational correction of off-targeting for CRISPR-Cas9 essentiality screens. Genomics; 2019. Available from: http://biorxiv.org/lookup/doi/10.1101/809970.

53. Murai J, Huang SyN, Das BB, Renaud A, Zhang Y, Doroshow JH, et al. Trapping of PARP1 and PARP2 by Clinical PARP Inhibitors. Cancer Research. 2012 Nov;72(21):5588–99. Available from: https://aacrjournals.org/cancerres/article/72/21/5588/576072/Trapping-of-PARP1-and-PARP2-by-Clinical-PARP.

54. Knezevic CE, Wright G, Remsing Rix LL, Kim W, Kuenzi BM, Luo Y, et al. Proteome-wide Profiling of Clinical PARP Inhibitors Reveals Compound-Specific Secondary Targets. Cell Chemical Biology. 2016 Dec;23(12):1490–503. Available from: https://linkinghub.elsevier.com/retrieve/pii/S2451945616303890.

55. Van Der Heijden FLAM, Weijers SA, Bleijerveld O, Kliza KW, Vermeulen M, Filippov DV. Proteome-Wide Profiling of Olaparib Interactors Using a Biotinylated Photoaffinity Probe. ChemBioChem. 2025 Mar;26(6):e202400882. Available from: https://chemistry-europe.onlinelibrary.wiley.com/doi/10.1002/cbic.202400882.

56. Ceballos-Arroyo AM, Munnangi M, Sun J, Zhang K, McInerney J, Wallace BC, et al. Open (Clinical) LLMs are Sensitive to Instruction Phrasings. In: Proceedings of the 23rd Workshop on Biomedical Natural Language Processing. Bangkok, Thailand: Association for Computational Linguistics; 2024. p. 50-71. Available from: https://aclanthology.org/2024.bionlp-1.5.

57. Chang YC, Huang MS, Huang YH, Lin YH. The influence of prompt engineering on large language models for protein–protein interaction identification in biomedical literature. Scientific Reports. 2025 May;15(1):15493. Available from: https://www.nature.com/articles/s41598-025-99290-4.

58. Kuligin L, Lammert J, Ostapenko A, Bressem K, Boeker M, Tschochohei M. Prompt design for medical question answering with Large Language Models. Machine Learning with Applications. 2025 Dec;22:100758. Available from: https://linkinghub.elsevier.com/retrieve/pii/S2666827025001410.

59. Deng C, Zhao Y, Tang X, Gerstein M, Cohan A. Investigating Data Contamination in Modern Benchmarks for Large Language Models. arXiv; 2023. Version Number: 2. Available from: https://arxiv.org/abs/2311.09783.

60. Xu C, Guan S, Greene D, Kechadi MT. Benchmark Data Contamination of Large Language Models: A Survey. arXiv; 2024. Version Number: 1. Available from: https://arxiv.org/abs/2406.04244.

61. The Gene Ontology Consortium, Aleksander SA, Balhoff JP, Carbon S, Cherry JM, Ebert D, et al. The Gene Ontology knowledgebase in 2026. Nucleic Acids Research. 2026 Jan;54(D1):D1779-92. Available from: https://academic.oup.com/nar/article/54/D1/D1779/8383826.

62. Ashburner M, Ball CA, Blake JA, Botstein D, Butler H, Cherry JM, et al. Gene Ontology: tool for the unification of biology. Nature Genetics. 2000 May;25(1):25–9. Available from: https://www.nature.com/articles/ng0500_25.

63. Zhao Y, Yu L, Wu X, Li H, Coombes KR, Au KF, et al. CEDA: integrating gene expression data with CRISPR-pooled screen data identifies essential genes with higher expression. Bioinformatics. 2022 Nov;38(23):5245–52. Available from: https://academic.oup.com/bioinformatics/article/38/23/5245/6762075.

64. Wolf T, Debut L, Sanh V, Chaumond J, Delangue C, Moi A, et al.. HuggingFace’s Transformers: State-of-the-art Natural Language Processing. arXiv; 2019. Version Number: 5. Available from: https://arxiv.org/abs/1910.03771.

65. for AI Innovation (DCAI) DC. Gefion;. Available from: https://dcai.dk/gefion.

