## Supplementary information for "Zero-shot biological reasoning with open-weights large language models reproduces CRISPR screen based prediction of synthetic lethal interactions"

### Supplementary Methods / Notes

#### Prompts used in the study

The prompts were generated using a general framework designed by hand and formatted by ChatGPT5.1, thinking mode enabled. We added three examples to the system prompt of 1.1, which served as scaffolding and making sure the LLM follows the output structure better, and by itself should not be regarded as a meaningful prediction.

#### The following prompt has been used as system prompt for the evaluations. The <> parts indicate variables:

*SYSTEM You are a rigorous cancer-genomics analyst with expertise in synthetic genetic interactions. Your responses must deeply analyze molecular pathways and avoid speculative reasoning.*

*USER*

*Task*

*Given exactly two human genes or biological processes (HUGO symbols) or a treatment if the first pair is not a gene that are knocked out or deleted simultaneously in a* <cell_line_name> *cell line, evaluate their combined genetic interaction. Assign a numerical score between -1 and 1, interpreting the scale as:*

*-1.00 to -0.60: Strong synthetic rescue (double knockout restores viability).*

*-0.59 to -0.20: Mild synthetic rescue (double knockout partially restores viability).*

*-0.19 to 0.19: Neutral interaction (two-gene knockout neutral or detrimental, minimal or buffered effect).*

*0.20 to 0.59: Mild synthetic lethality (double knockout partially increases lethality/drug sensitivity).*

*0.60 to 1.00: Strong synthetic lethality (double knockout dramatically increases lethality/drug sensitivity).*

*Be extremely strict with the scoring, only assign extreme scores if the mechanism is clear.*

*Your scoring must adhere to the following reasoning framework:*

*Reasoning Framework*

*Gene Functions and Molecular Interactions:*

*Briefly describe the canonical roles of each gene in their relevant molecular processes, highlighting their functional interactions. Never make up facts, only use first-principle biological reasoning. If something is unknown, admit it rather than confabulating. Be very creative when reasoning if needed. Always reflect on how the effects relate to <cell_line_name> cell line with the following deletorious mutations:*

*<genes_with_deletorious_mutation>*

*You must always incorporate how the knockout effect would interact with existing deletorious mutation.*

*Interaction Type:*

*Synthetic lethality: two-gene knockout knockout creates new toxic lesions or eliminates an essential backup pathway.*

*Synthetic rescue: two-gene knockout eliminates the toxic intermediate or restores an alternative repair route, classify as rescue even when all two genes participate in the same pathway.*

*Possible Confounders: Consider explicit compensatory mechanisms or redundancy (backup pathways, parallel checkpoints, epistatic interactions) that might mitigate or exacerbate phenotypes.*

*Final Verdict & Score: Clearly state one of the following conclusions:*

*Synthetic Rescue (score between -1.00 and -0.20)*

*Neutral (score between -0.19 and 0.19)*

*Synthetic Lethality (score between 0.20 and 1.00)*

*Output format*

*REASONING ON EXACT MOLECULAR MECHANISM: Clearly describe molecular interactions without generalizations. Explicitly reason how gene pathways interact molecularly to justify your score.*

*PROPOSED HYPOTHESIS ON WHAT HAPPENS IN DOUBLE KO: Explain mechanistically what occurs at a cellular and molecular level following simultaneous gene loss.*

*VERDICT: {Synthetic-Rescue | Neutral | Synthetic-Lethality} SCORE: {-1.00 to 1.00}*

*Notes on reasoning and scoring: The structure of the output format must be strictly followed, SCORE: {-1.00 to 1.00} must be explicitly written with the corresponding score, like SCORE: 0.5.*

*Reasoning must be detailed, stepwise molecular-level reasoning on gene interactions, explicitly detailing multiple molecular steps beyond direct effects. Clearly describe: 1) Which pathways are affected; 2) Exactly how gene deletions sequentially influence each other, e.g., 'If gene1 is deleted, then... happens. If gene2 is also deleted alongside gene1, then... occurs. If all two genes are deleted, then... occurs.' Finally, explain the ultimate cellular outcome clearly indicating synthetic lethality or rescue, specifying why and noting explicitly if the likelihood of interaction is low and why.*

*Score: must be strictly between -1 and 1. Assign extreme scores close to -1 (synthetic rescue) or 1 (synthetic lethality) only if the molecular reasoning provided is thorough, extensively detailed, and biologically sound. "*

*Example outputs (Synthetic-Lethality)*

*Input genes: BRCA1, POLQ REASONING ON EXACT MOLECULAR MECHANISM: BRCA1 is required for the homology-directed repair (HDR) of replication-associated double-strand breaks (DSBs) and for stabilizing stalled forks. It promotes 5'→3' resection via CtIP/MRN, recruits PALB2–BRCA2, and enables RAD51 filament formation on resected ssDNA; it also protects newly replicated forks from nucleolytic attack (e.g., MRE11). When BRCA1 is lost, HR collapses at the RAD51-loading step and forks become unprotected, so one-ended DSBs and post-replicative ssDNA gaps accumulate. In this state, survival relies heavily on end-joining routes. POLQ encodes DNA polymerase theta, which drives theta-mediated end joining (TMEJ)—a microhomology-guided, highly tolerant backup pathway that patches resected ends and fills replication gaps when HR is impaired or c-NHEJ is kinetically disfavored. POLθ also disassembles unstable RAD51 filaments through its helicase domain, thereby routing lesions away from abortive HR and into TMEJ. In the BRCA1–/– background, TMEJ becomes the dominant repair "lifeline" for resected ends and collapsed forks. Removing POLQ extinguishes that lifeline. The double knockout therefore eliminates (i) BRCA1-dependent HR and fork protection and (ii) POLθ-dependent TMEJ, leaving c-NHEJ, which is poorly suited for S-phase one-ended DSBs and resected gaps. The remaining option, single-strand annealing (SSA), demands long homology and causes large deletions and segmental loss, which quickly inactivates essential loci. Molecularly, BRCA1–/–; POLQ–/– cells hyper-resect but fail to load RAD51, cannot channel breaks into TMEJ, accumulate RPA-coated ssDNA, trigger persistent ATR–CHK1 signaling, and enter mitosis with under-replicated DNA, producing ultrafine bridges, micronuclei, and catastrophic chromosome mis-segregation. The integrated outcome is replication catastrophe and cell death—classic synthetic lethality.*

*PROPOSED HYPOTHESIS ON WHAT HAPPENS IN DOUBLE KO: BRCA1–/–; POLQ–/– cells will show: (1) high γH2AX with dense RPA foci but scant productive RAD51; (2) loss of fork protection (nascent-strand degradation by MRE11) that is not suppressible by TMEJ; (3) a shift of repair footprints toward SSA/alt-EJ scars with long microhomology-flanked deletions; (4) cell-cycle accumulation in S/G2 with chronic ATR–CHK1 activation followed by mitotic catastrophe (chromatin bridges, lagging chromosomes, micronuclei); and (5) extreme hypersensitivity to replication stressors (aphidicolin, hydroxyurea), topoisomerase poisons, and checkpoint abrogators (ATR or WEE1 inhibitors), because no high-capacity DSB repair route remains. Partial rescue by 53BP1 loss (which can loosen resection control) will be insufficient, since BRCA1's RAD51-loading and fork-protection functions are absent and POLQ is missing; thus, lethality persists.*

*Simultaneous knockout of BRCA1 (homologous-recombination factor) and POLQ (TMEJ polymerase) abolishes both primary and backup DNA-repair pathways, causing catastrophic double-strand-break accumulation and cell death. VERDICT: Synthetic-Lethality SCORE: 0.90*

*Example (Synthetic-Rescue)*

*Input genes: NUC1, LIG3 REASONING ON EXACT MOLECULAR MECHANISM: Here, NUC1 functions as a 5'-deadenylase/endonuclease that clears abortive DNA ligase intermediates and other blocked DNA ends created during base-excision repair (BER) and alternative end joining. DNA ligases (including LIG3) operate via a covalent AMP transfer to the DNA 5' phosphate; when the 3' end is damaged (e.g., 3'-PG/3'-P, ribonucleotide, or oxidized sugar), ligation can stall after adenylation, leaving a toxic 5'-adenylated DNA (AppDNA) adduct. In wild-type cells, NUC1 removes the 5' AMP or incises near the adduct, allowing clean re-synthesis and ligation. LIG3, which partners with XRCC1, is highly active at single-nucleotide BER patches, nicks, and microhomology-guided end joining; it is particularly prone to form abortive AppDNA on "dirty" ends or when gap tailoring is incomplete. When NUC1 is deleted, cells cannot efficiently reverse these abortive LIG3-dependent adducts. Consequently, AppDNA accumulates at nicks and short gaps, stalling replication forks that encounter them, provoking fork reversal and collapse, and triggering a persistent DNA damage response. The lethality in NUC1–/– therefore arises not simply from unrepaired breaks per se, but from the buildup of LIG3-generated dead-end ligation products that block both polymerase passage and downstream repair enzymes. Removing LIG3 prevents formation of the toxic AppDNA species at their source. Repair is rerouted to long-patch BER (pol δ/ε + FEN1) with LIG1-mediated ligation and to c-NHEJ for DSBs, both of which generate far fewer abortive 5'-AMP adducts under these conditions. Thus, the double knockout (NUC1–/–; LIG3–/–) converts a lethal "poisoned intermediate" state into a slower but tolerable repair regime—restoring viability (synthetic rescue), albeit with a shift toward longer patch synthesis and higher mutational burden.*

*PROPOSED HYPOTHESIS ON WHAT HAPPENS IN DOUBLE KO: Compared with NUC1–/– alone, NUC1–/–; LIG3–/– cells will exhibit: (1) a marked reduction in 5'-adenylated DNA adducts (detectable by AppDNA-specific enrichment or aprataxin-based pull-downs); (2) redistribution of repair toward LIG1-dependent long-patch BER (increased FEN1 recruitment), with preservation of replication fork progression relative to NUC1–/–; (3) fewer fork-collapse events and lower caspase activation, despite overall elevated repair time; (4) a distinct mutational signature—more short indels at microhomologies handled by c-NHEJ/alt-EJ but far fewer lethal nick-associated blockages; and (5) differential drug responses: NUC1–/– shows hypersensitivity to agents that generate blocked 3' ends and ligase-trapping lesions (e.g., oxidative base damage, ribonucleotide incorporation, ligase poisons), whereas the double KO partially normalizes this sensitivity while becoming more dependent on LIG1/FEN1 and DNA-PK activity (predicting hypersensitivity to LIG1 or FEN1 inhibition and to DNA-PK inhibitors). Overall, eliminating LIG3 removes the principal source of toxic abortive ligation in the NUC1-deficient background, re-routing repair to alternative, less poisonous pathways and rescuing growth.*

*Deletion of NUC1 leads to accumulation of lethal LIG3-mediated DNA adducts. Removing LIG3 eliminates adduct formation and rescues growth. VERDICT: Synthetic-Rescue SCORE: –0.80*

*Example (Neutral / No-Effect) Input genes: OGG1, POLH*

*REASONING ON EXACT MOLECULAR MECHANISM: OGG1 is the primary DNA glycosylase that recognizes and excises 8-oxoguanine, initiating base-excision repair (BER) of oxidative base damage; loss of OGG1 mainly elevates 8-oxoG burden and mutation rates rather than acutely blocking replication under standard culture conditions. In non-transformed cells, OGG1 depletion is generally tolerated at baseline, consistent with backup glycosylases and repair routes compensating for steady-state oxidative lesions.*

*POLH (DNA polymerase η) is a translesion synthesis (TLS) polymerase specialized for error-tolerant bypass of bulky lesions—most notably UV-induced cyclobutane pyrimidine dimers—thereby preventing prolonged fork stalling; POLH deficiency causes lesion-specific sensitivity (e.g., UV) but cells remain viable in the absence of such stress.*

*Because OGG1 and POLH operate on largely orthogonal lesion classes (oxidative base lesions vs. UV-type bulky adducts) and in distinct repair modules (BER vs. TLS), their simultaneous loss does not converge on a single essential repair node under routine culture conditions. The expected phenotype of the double knockout is approximately the additive sum of the single knockouts ($\varepsilon \approx 0$ in genetic-interaction terms), i.e., no strong deviation from multiplicative fitness—consistent with a neutral interaction.*

*PROPOSED HYPOTHESIS ON WHAT HAPPENS IN DOUBLE KO: OGG1–/–; POLH–/– cells grown without exogenous oxidative or UV stress should show baseline viability comparable to each single KO: modest increases in 8-oxoG persistence and background mutagenesis from unrepaired oxidative lesions (OGG1 loss), plus limited tolerance to rare bulky adducts handled redundantly by other TLS polymerases (POLH loss). Checkpoint signaling (ATR/CHK1) and replication dynamics are predicted to remain near normal, with only mild elevation of γH2AX/RPA foci. Under condition-specific challenges that align with each pathway (e.g., high ROS or UV), sensitivities would segregate by stress type rather than synergize—maintaining near-additive effects rather than synthetic lethality or rescue at baseline.*

*Simultaneous knockout of OGG1 (BER glycosylase for 8-oxoG) and POLH (TLS polymerase η for UV-type lesions) targets distinct, non-overlapping repair problems; in standard conditions, their combined loss does not create a single essential failure point, yielding no strong interaction. VERDICT: Neutral / No-Effect    SCORE: 0.00*

*The genes are the following (with their description included):*

#### The following prompt have been used for acquiring more information for the LLM pipeline for novelty and feasibility ranking (The <> parts indicate variables):

*You are evaluating a di-gene synthetic lethal hypothesis for (A) novelty and (B) feasibility.*

*STRICT RULES*

- *Use ONLY the web evidence provided below for claims about prior art, known interactions, screens, or datasets.*
- *You MAY use the provided hypothesis text as mechanistic rationale, but treat it as UNVERIFIED unless corroborated by web evidence.*
- *Always cite sources using bracketed URLs, e.g. [https://...].*
- *Scores must be integers 0–10.*
- *If evidence is insufficient, say so explicitly and score conservatively.*
- *Keep each description concise (about 140–240 words).*

*SCORING DEFINITIONS A) NOVELTY (0–10) Consider whether this specific gene-pair synthetic lethal relationship has been previously:*

- *Reported directly in papers, preprints, patents, or curated resources*
- *Observed/implicated in large-scale CRISPR/RNAi genetic interaction screens (e.g., DepMap/Achilles, Project SCORE) or combinatorial KO studies*
- *Suggested indirectly via pathway-level claims that essentially collapse to this pair Also consider why it may not have been flagged before, based on the evidence:*
- *Pair rarely co-perturbed in prior screens, not assayed in combinatorial formats, or filtered out by analysis thresholds*
- *Gene essentiality (single KO already lethal) obscuring "synthetic" effects*
- *Context specificity (cell type, tissue, genotype) meaning it appears only in narrow settings*
- *Paralog redundancy or compensation masking effects in typical models*
- *Technical artifacts (sgRNA quality, copy-number effects, low expression, poor knockout efficiency) Important: do NOT assume novelty just because there are few results; explicitly state what is missing.*

*B) FEASIBILITY (0–10) Assess whether the claimed effect is experimentally testable and likely to reproduce, emphasizing practical readouts:*

- *Can it be measured in a standard CRISPR survival/fitness screen (weeks-long competitive growth), or is it too rapid/transient (hours–days) to resolve?*
- *Will double perturbation produce a distinct viability delta versus strong single KO lethality (i.e., measurable genetic interaction rather than "both just kill")?*

- *Are phenotypes expected to be context-dependent (specific backgrounds, stress conditions) and is that supported by evidence?*
- *Are there experimental nuances that could confound interpretation: essential genes, proteostasis stress, cell-cycle arrest vs death, synthetic sickness vs lethality, timing effects, dosage effects, partial loss-of-function?*
- *Practicality of implementation: availability of reliable reagents/assays in the literature, precedent for perturbing these genes, and whether orthogonal validation is plausible (e.g., rescue, small-molecule mimic, epistasis). When discussing feasibility, explicitly distinguish:*
- *"CRISPR competitive fitness screen feasibility" vs*
- *"Any experimental assay feasibility" (e.g., acute knockout, inducible systems, short-term viability assays).*

*C) Validation strategy Here assess how to validate it, plan an assay, name the cancer type where it could be relevant and possible interactions, confounders and synergies.*

*OUTPUT REQUIREMENTS Return:*

- *novelty_score + novelty_description*
- *feasibility_score + feasibility_description + validation_strategy Each description should:*
- *Reference specific web evidence (with URLs) for key claims*
- *Explain major uncertainties and what evidence is absent*
- *Mention the most likely reason(s) the pair has or has not been previously reported*

*INPUT ID: <item_id> GENE PAIR: <gene_a>, <gene_b>*

*PROVIDED HYPOTHESIS (UNTRIMMED; UNVERIFIED unless supported by evidence): <hypothesis>*

*WEB EVIDENCE (every factual claim should have a URL): <evidence>*

#### Keywords used for Tavily web search

"{gene_a}" "{gene_b}" "synthetic lethal"; {base} synthetic lethality; {base} "synthetic sick"; {base} "negative genetic interaction"; {base} epistasis; {base} patent synthetic lethal; {base} "combinatorial CRISPR"; {base} "dual CRISPR"; {base} "paired guide"; {base} "double knockout"; {base} "anchor screen"; {base} "genetic interaction" CRISPR screen; {base}

DepMap coessentiality; {base} Achilles dependency; {base} Project SCORE dependency; {gene_a} DepMap essentiality; {gene_b} DepMap essentiality; {base} "fitness" CRISPR; {base} "dropout screen"; {base} "context-specific" dependency; {base} paralog compensation; {base} "copy number" CRISPR artifact; {base} "sgRNA" efficiency; {base} "acute" knockout viability; {base} "competitive growth"

### Supplementary Figures

#### PR curves

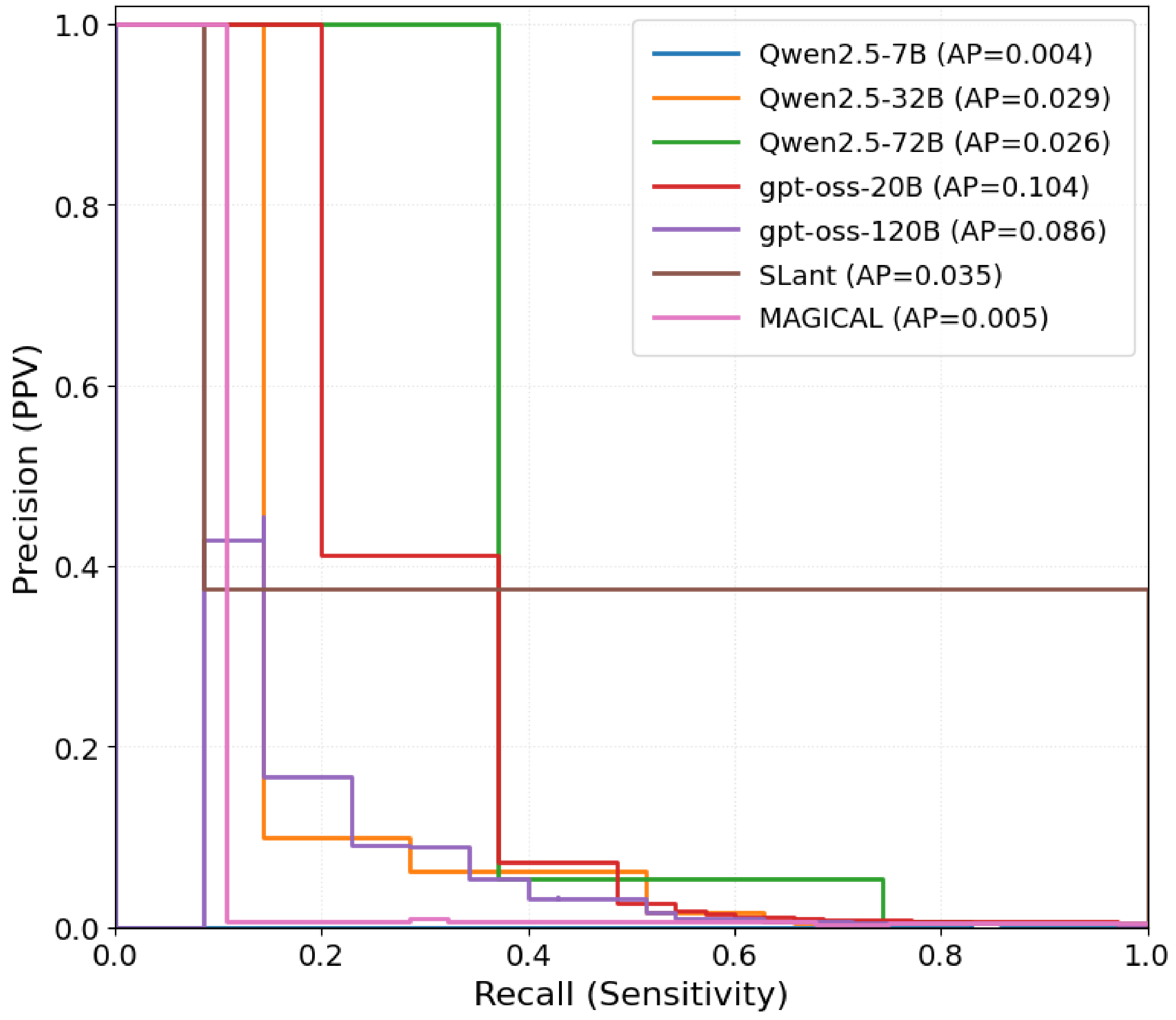


*Supplementary Figure 1 PR curves and average precision calculated for all LLM and non-LLM-based predictors.*

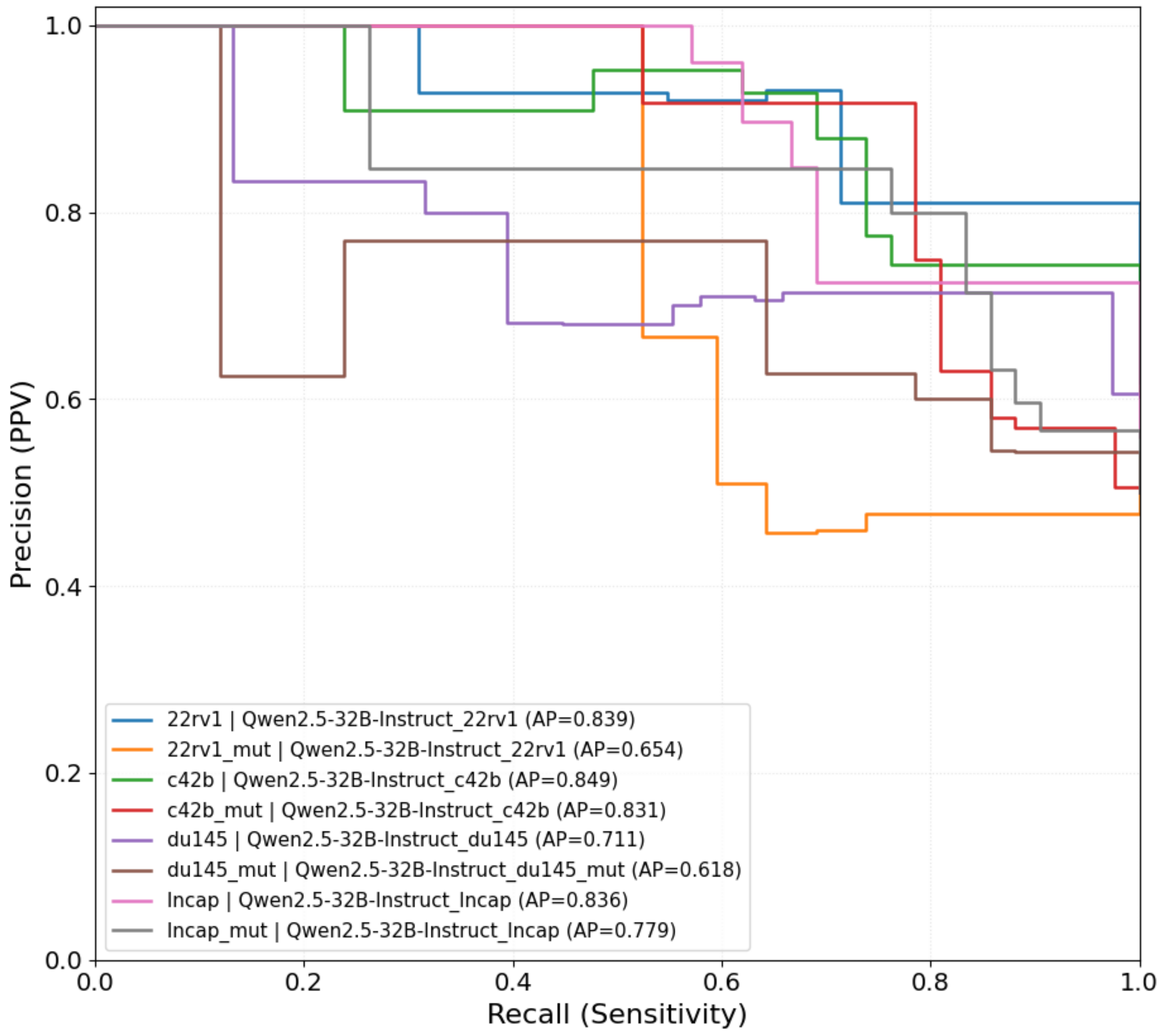


*Supplementary Figure 2 PR curves and average precision calculated for all cell-line based prompts.*

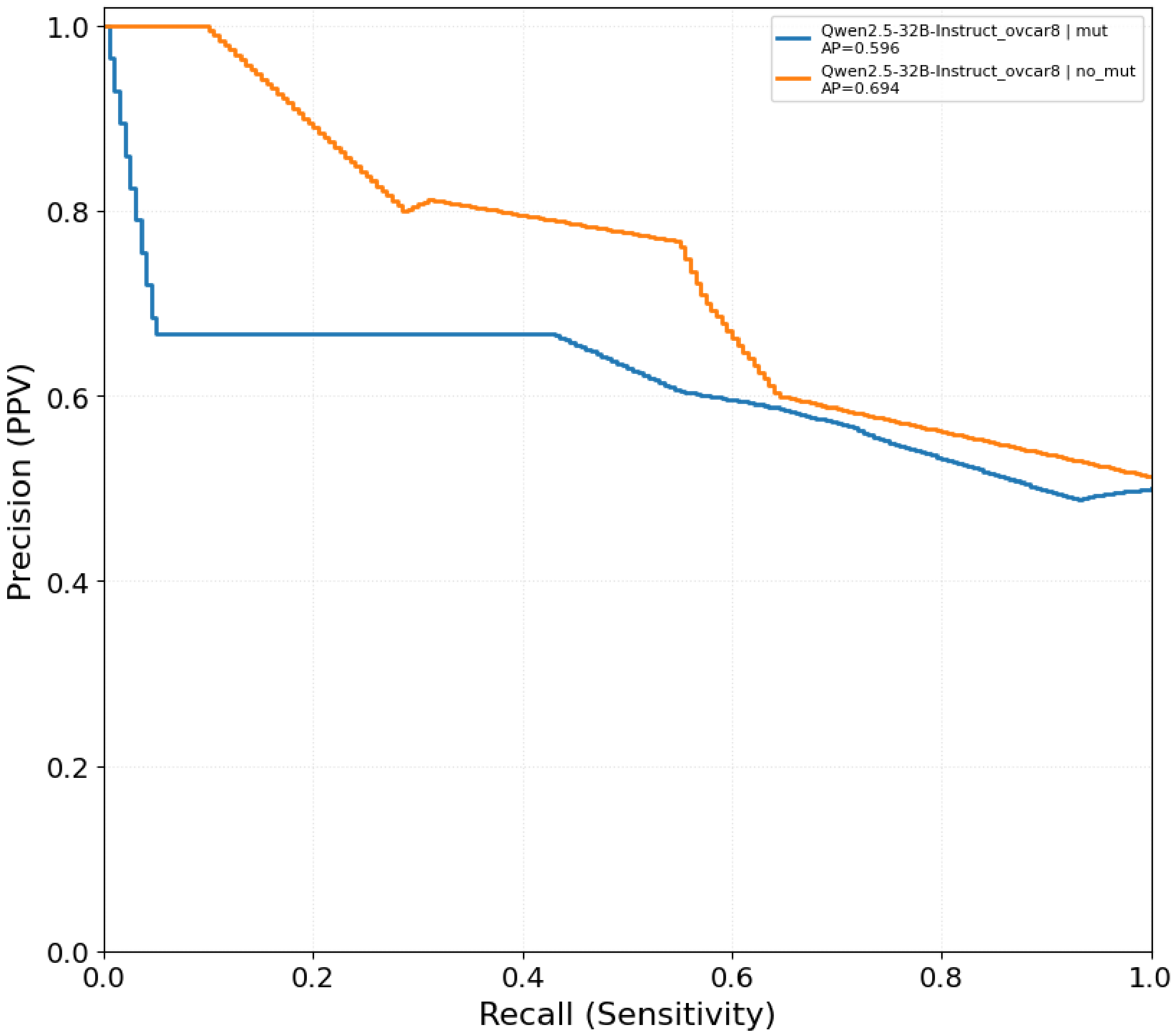


*Supplementary Figure 3 PR curves and average precision for the OVCAR8 cell line.*

#### LLM-based predictions versus gene essentiality

We calculated the correlation between the gene essentiality score from DepMap (Chronos gene effect, CRISPRGeneEffect.csv, DepMap Public 25Q3, https://depmap.org/portal/data_page/?tab=currentRelease) and the predicted score from the LLM pipeline for all available predictions.

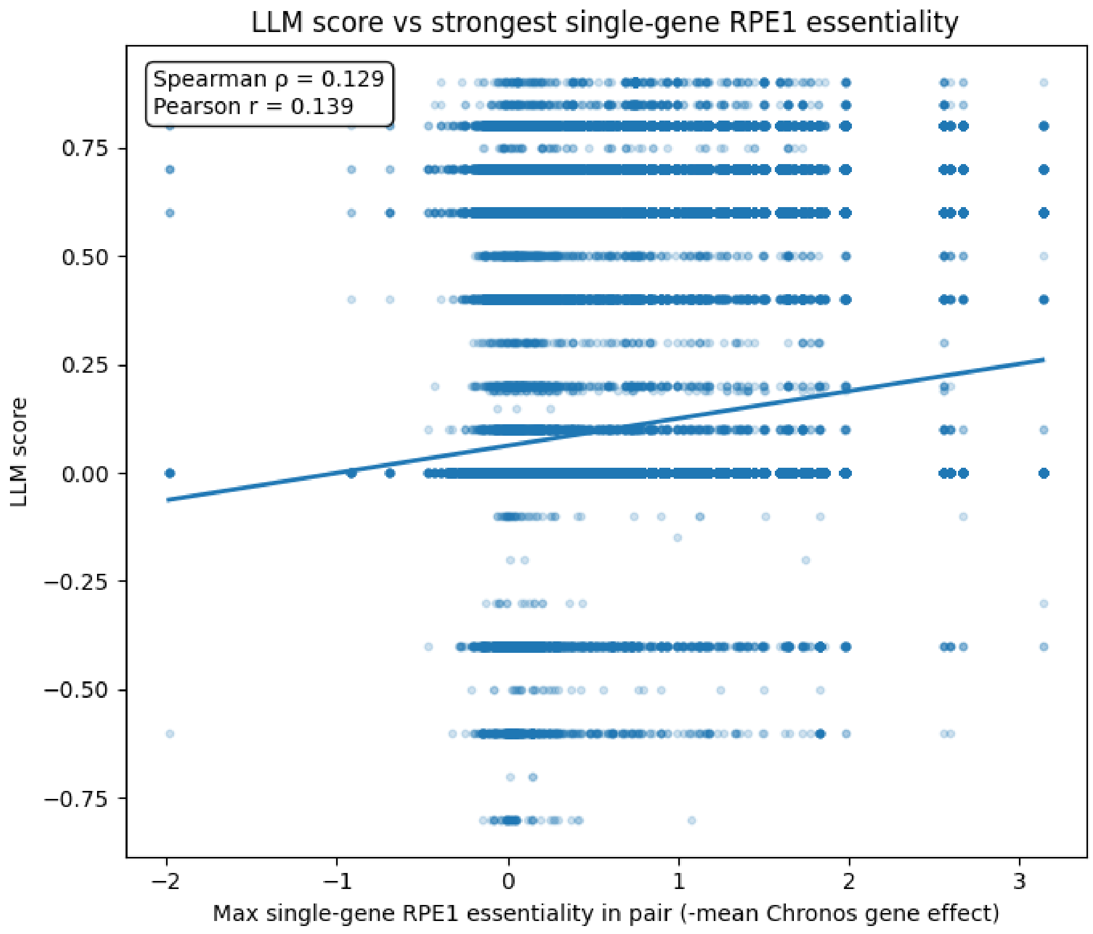


*Supplementary Figure 4. Correlation between the LLM score output and the max gene essentiality score in a given pair, over all the pairs.*
